# Stimulus-response correlation analysis dissociates spatiotemporal cortical networks supporting speech production

**DOI:** 10.64898/2026.05.30.729015

**Authors:** Arka N. Mallela, Raouf Belkhir, Eliza Reedy, Hussam Abou-Al-Shaar, Jiahao Chen, Julien Dirani, Bradford Z. Mahon, Jorge A. González-Martínez

**Author notes:** Corresponding author: Jorge A. González-Martínez MD PhD, 200 Lothrop Street, Suite B-400, Pittsburgh, PA 15213. Co-senior authors (Bradford Z. Mahon, Jorge A. González-Martínez).

## Abstract

**Introduction:** Understanding the spatiotemporal distribution of cortical activation during language production is a central question in cognitive neuroscience with broad clinical applications. High spatial/temporal resolution recording over multiple brain regions and specific psycholinguistic manipulations with testable behavioral predictions are necessary to separate neural variance attributable to processing stages.

**Objective:** We combine a delayed naming paradigm with intracranial electrophysiology to identify spatiotemporally dissociated cortical networks supporting stages of speech production.

**Methods:** Subjects (20 healthy, 15 intracranial electrophysiology) are presented visual stimuli to name but are instructed to delay response until a go cue at 0/400/1000ms. Exploiting the temporal variance induced by delay, we calculate stimulus/response-locked correlation of high gamma (70-200Hz) activity at each contact. We compare shifts in stimulus-response correlation space to response time (RT) to validate separation of networks.

**Results:** Behaviorally, the factor delay modulated the effect of lexical selection on RT (p=0.021) but did not modulate the effect of phonological planning (p=0.4), confirming the paradigm separates processing stages. Stimulus-response correlation analyses revealed globally-invariant structure across subjects. Certain regions showed strong stimulus-locking (occipital lobe) or response-locking (motor cortex), but most regions were intermediate. Shifts in correlation space identified dissociated networks that predicted substantial variance in RT from lexical selection (R^2^=0.64) and phonological density (R^2^=0.75). Using a leave-one-subject-out cross validation approach, these shifts explained 31.4% of variance in RT at the trial level.

**Conclusion:** Stimulus-response correlation space reveals stable spatiotemporal signatures of psycholinguistic processing that generalize across individuals. This approach permits hyperalignment of neurophysiological data and functional separation of cortical networks in speech production across subjects.

## Introduction

The spatiotemporal distribution of cortical activation during language production is a central question in cognitive neuroscience with immense implications for understanding normal speech function and the pathophysiology of aphasia and deficits following neurosurgery^1,2^. In the context of picture naming, current leading models^2–4^ suggest a largely sequential model of activation with initial activity in occipito-temporal regions, followed by lexical access in the middle temporal gyrus (MTG), phonetic encoding in MTG and superior temporal gyrus (STG), syllable segmentation in the left inferior frontal gyrus (IFG, including the classical “Broca’s area”) and finally motor programming in speech motor cortex (SMC)^1^. This general model has found broad support through a variety of modalities including scalp EEG^5–7^, functional MRI^8–12^, and intracranial EEG^13^, particularly electrocorticography with subdural grid electrodes^14,15^ and direct cortical stimulation^16–21^. The degree to which this system is strictly hierarchical and ordered as opposed to cascading and interactive has been debated^22–25^.

A fundamental challenging to answering this question is finding a method for measuring brain activity with 3 properties – high temporal resolution, high spatial resolution, and simultaneous recording across multiple brain regions. Scalp EEG and MEG provide high temporal resolution, but the signal is an amalgamation from thousands of cortical sources per channel^26^. FMRI provides spatial resolution but is unable to distinguish activation at the millisecond level. Intracranial recording of the local field potential (LFP) addresses the issues of spatial and temporal resolution but is ultimately limited by the coverage of the intracranial electrodes. Subdural grids and intracranial stimulation are often limited to the dorsolateral surface of the hemisphere with limited coverage of sulcal and medial structures. Stereo EEG (SEEG) is invasive monitoring methodology to investigate drug-resistant epilepsy in which one or more anatomo-electrical clinical hypotheses are interrogated with stereotactic placement of depth electrodes^27–30^. This distributed monitoring permits simultaneous recording of multiple cortical regions, including the depths of sulci and the medial surface of the hemisphere, in exchange for less dense coverage in each region. Thus, we hypothesized that distributed, simultaneous recording at high spatial and temporal resolution could address questions regarding the spatiotemporal distribution of the stages of language production.

Here, we rely on well-described psycholinguistic manipulations of stimulus type^2,31,32^, lexical frequency^22–24^,, phonological density^33,34^ that interface with particular psycholinguistic processes, namely visual recognition, lexical access, and phonological planning, respectively^1^. We separate stimulus presentation with a variable delay period (0, 400, 1000ms) to separate processes involved in language planning and speech production. We first test this hypothesis behaviorally in both a cohort of 20 healthy subjects and 15 patients who underwent intracranial monitoring with SEEG for drug-resistant epilepsy. Subsequently, we interrogate the spatiotemporal implementation of these processes in the cortex. To compare electrophysiological data across subjects and cortical regions, we develop a novel correlation space analysis that permits simultaneous analysis of cortical networks that leverages the behavioral temporal variance created by the delay. Finally, we demonstrate that the activity in these networks predicts the behavioral effects that we observed, tying the neurophysiology to the psycholinguistic model of speech. These results support a partially sequential model but with considerable distributed and interactive cortical activity.

## Methods

### Subject selection

This study was approved the Institutional Review Boards at the University of Pittsburgh (intracranial electrophysiology subjects) and Carnegie Mellon University (healthy controls). For healthy controls, 20 subjects were recruited from undergraduate/graduate students to participate in behavioral testing. Intracranial electrophysiology subjects (N=15, 40% female, age 34 ± 13 years old) were recruited from the practice of the senior author. Intracranial subjects were patients with drug-resistant epilepsy who underwent invasive electrophysiological monitoring with stereo EEG and were tested during their stay in the epilepsy monitoring unit. All subjects provided informed consent. For both populations inclusion/exclusion criteria were age over 18, right-handed, English as a first language, and lack of any speech or language disorder. Additional criteria for intracranial subjects were lack of any major anatomic abnormalities, major prior resective brain surgery, lack of post-operative hemorrhage, and ability to complete the task.

### Delayed naming paradigm

Stimulus presentation was performed with PsychoPy^35^ using a custom built workstation for healthy subjects and for use in the epilepsy monitoring unit. Each trial consisted of a 500ms fixation cross, followed by a 250ms blank screen, after which stimulus was presented either with the go cue (green outline) immediately or after a delay period of 400ms or 1000ms. In delayed trials, the stimulus was initially outlined with a red square. Stimuli included high/low frequency pictures from the Snodgrass task^36^, and words and pseudowords drawn from the PALPA word/nonword repetition tasks^37^. Estimates of phonological density were calculated using the IPhOD database^38^. Audio transcripts of subject responses was collected using a directional microphone and scored offline for accuracy, response time after go cue (RT) and response duration.

### Behavioral analysis

All subjects were examined for accuracy and excluded if overall accuracy was less than 90%. No subject was excluded by this criterion. We used linear mixed effects modeling of response time, with subject as a random effect, against the desired psycholinguistic variables (e.g. stimulus type, picture lexical frequency, repetition, phonological density, delay). We investigated the interaction of delay and other variables using Type III ANOVA using Satterthwaite’s method. Healthy controls and intracranial subjects were analyzed separately.

### Intracranial electrophysiology data acquisition and preprocessing

Intracranial subjects underwent stereotactic implantation of stereo EEG using DIXI Medical (Marchaux-Chaudefontaine, France) Microdeep electrodes (platinum/iridium, 0.8mm diameter, 2mm contact size, 1.5mm spacing) with 8 - 18 contacts per electrode shaft. Electrode implantation patterns were determined solely by clinical need in all cases. A total of 2975 contacts across 15 subjects (198±41 contacts per patient) were implanted. 242 (8.1%) were excluded due to proximity to the seizure onset zone, spiking activity, or technical issue (broken contact, contact in skull, etc.).

Data was acquired on Natus Neuroworks Quantum system (Natus Medical Inc., Pleasanton, CA), a clinical-grade EEG acquisition platform at sampling rate of 2048 or 1024 Hz with a hardware bandpass filter of 0.01–3000 Hz, with high input impedance and low-noise differential amplifiers. A DC trigger was passed from the stimulus presentation computer to Natus to ensure accurate behavioral-electrophysiological synchronization. Data was referenced to a common average reference^39^, band pass filtered from 0.1-200 Hz, notch filtered at 60 Hz and higher harmonics and then epoched in stimulus or response-locked alignments. Epochs with excess noise were excluded.

Electrode position was confirmed with post-operative high-resolution CT (0.625mm^3^ isotropic resolution) and co-registered to preoperative T1-weighted MRI (MPRAGE, 1mm^3^ isotropic resolution). Contacts were segmented in native subject space using CURRY (Compumedics USA Inc, Charlotte, NC). Subject MRI scans were nonlinearly registered with tissue segmentation using SPM12^40^ to the MNI152 ICBM 2009c nonlinear asymmetric template^41^. Contact locations for all subjects were transformed into MNI space for subsequent analyses. The Harvard-Oxford atlas (max-prob25, 1mm^3^ resolution)^42^ was used for contact location analysis.

### Time frequency analysis

We utilized a similar time frequency methodology as other reports^43^. High gamma activity (70-200 Hz) was extracted analytic amplitude of the Hilbert transform following bandpass filtering with a two-pass (forward-backward) FIR filter. The amplitude was smoothed using the Savitzky-Golay filter (third order, 201ms frame length) and Z-scored to a pre-stimulus baseline of 500ms. Subsequently, the signal was downsampled to 100 Hz and the first and last 250ms of each epoch was trimmed to avoid edge effects. All electrophysiological preprocessing and time frequency analysis was performed with MNE Python^44^.

### Stimulus-response correlation space analysis

Prior to any subsequent analysis, electrode contacts were thresholded for response at any point in stimulus-locked epoching compared to pre-stimulus baseline using a t-test with a threshold of p < 0.05 with FDR. To compute stimulus/response correlation, for each contact in each subject, the smoothed, Z-scored, high gamma activity was correlated on a trial-to-trial basis for all trials for that subject and averaged over all pairs of trials at each contact in stimulus or response-locked epoching respectively. For investigations of the effects of psycholinguistic manipulations on stimulus-response correlation space, stimulus/response correlation was carried out over the subset of trials with each condition (e.g. high frequency pictures, low frequency pictures).

To classify electrodes in correlation space, electrode contacts with less than 0.025 shift in correlation space (norm of stimulus and response correlation shifts) were discarded, and remaining electrodes were classified on the basis of which quadrant in stimulus/response correlation change space they occupied.

To extract time course for each cluster, for each subject, the relevant epochs were filtered and the electrodes in each cluster were selected. The signal in these electrodes/epochs were averaged across all subjects.

### Volumetric analysis

As stereo EEG is a three-dimensional exploration of the brain, we utilized volumetric reconstruction for primary analyses and surface rendering for display purposes. For quantitative data (e.g. correlation per electrode), volumetric reconstruction of electrode data was performed by averaging all electrode values within 1cm of each voxel of the brain. For categorical data (e.g. classification), we used a K-nearest neighbor approach (5 neighbors within at most 1cm of voxel) at each voxel. Surface rendering over the fsaverage inflated template was performed and displays voxels within 3mm of the cortical surface using the python package *nilearn*.

### Response time vs. stimulus-response correlation space analysis

For each psycholinguistic variable, the mean shift in stimulus correlation and response correlation was computed for each cluster for each subject. Change in stimulus correlation and response correlation for each cluster were linearly regressed, separately then together, against mean change in response time per subject, treating each subject as an independent observation.

Subsequently, shifts in stimulus-response correlation space for all clusters of interests (corresponding to psycholinguistic variables/processes) were regressed against trial level RT. For each trial, we computed the cluster shift for each relevant process (e.g. stimulus type, lexical frequency, phonological density) according to the direction of manipulation (e.g. high vs. low) in that subject, providing a subject and trial specific estimate of the shifts in correlation space for each of our psycholinguistic manipulations. We then regressed this against trial-level RT. We used an elastic net linear regression with L1 and L2 regularization (regularization parameter = 2, L1 ratio = 1) to account for multicollinearity, feature selection, and overfitting, with leave-one-subject-out cross validation to determine the variance in RT explained (R^2^) by shifts in correlation space.

### Statistical analysis

Behavioral statistical analysis was performed using R (The R Foundation for Statistical Computing, Vienna, Austria). Regularized linear regression was performed using *scikit-learn*^45^ in Python 3.12. P < 0.05 was the preset threshold for significance for all analyses.

### Data availability

Data, including deidentified neuroimaging, electrophysiological, and response time data, is available upon reasonable request to the first or senior authors.

## Results

### Delayed naming paradigm

To temporally separate out processes involved in speech planning during picture naming, e.g. visual recognition, lexical selection, from those involved in production, e.g. syllabification and phonological encoding, we developed a delayed naming paradigm. Subjects were instructed that they would be shown visual stimuli but to not produce the response until a go cue (green square around stimulus) was presented. In the no (0 s) delay condition, subjects immediately received the go cue. In the 400 ms and 1000 ms conditions, the stimulus was surrounded by a red square which became green after the delay (**Figure 1A**). Stimuli consisted of high and low frequency pictures (Snodgrass items^36^), printed words, and printed pseudowords (**Figure 1B**). Subjects completed 4 blocks of 75 items each (300 total) with up to 3 repetitions of stimuli. Healthy subjects (N=20) completed 6000 trials, while intracranial electrophysiology subjects (N=15) completed 4113 trials with overall 94.8% accuracy.

**Figure 1:**
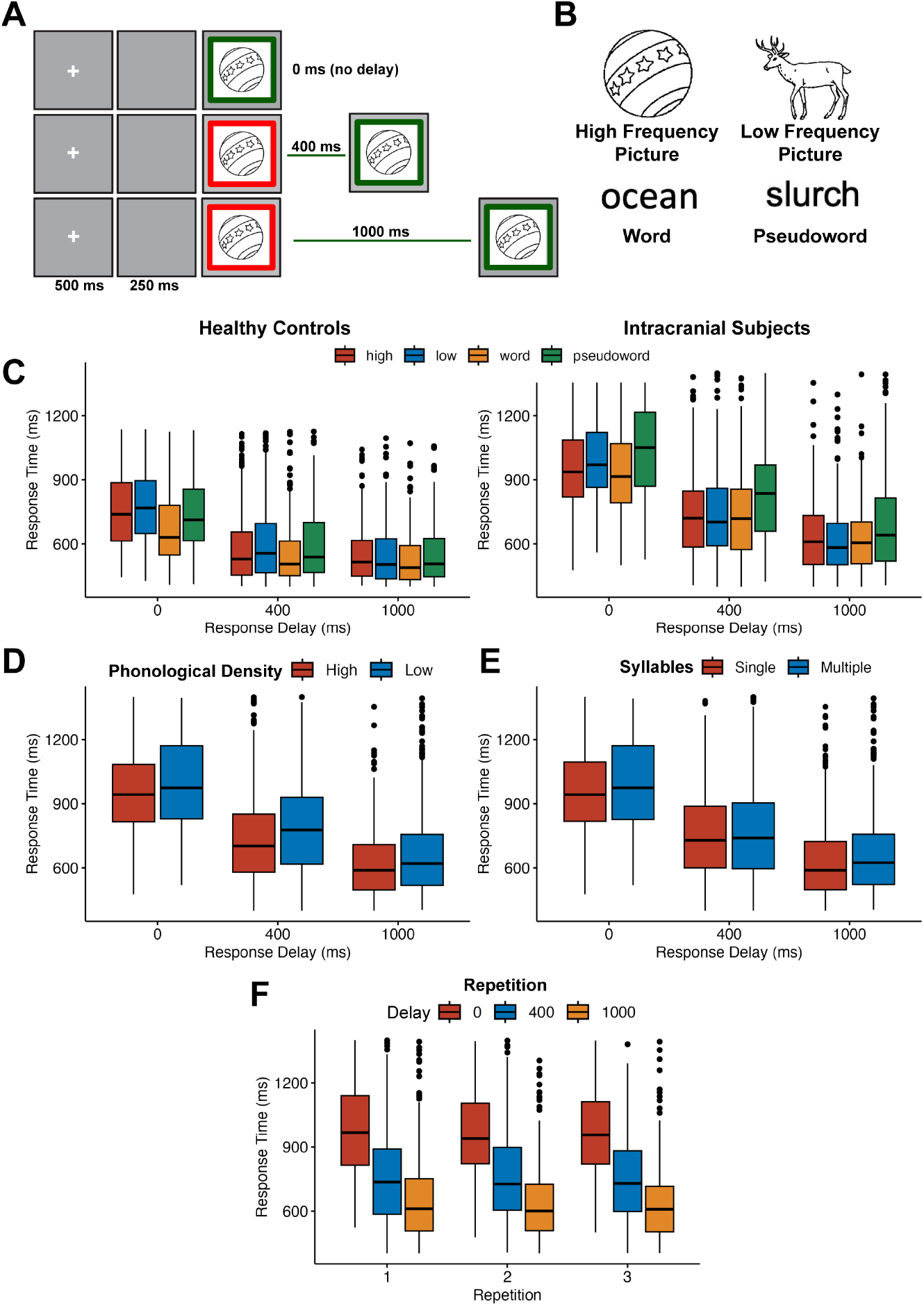
Delayed naming paradigm and behavioral results. **A**: Subjects are presented visual stimuli with a variable delay (0, 400, 1000ms) between stimulus presentation and response cue. Each trial begins with a 500ms fixation cross, followed by 250ms blank screen. The stimulus is then presented, either with a green outline, cuing immediate production or a red outline, indicating that the subject should prepare but not produce the response. In the 400/1000ms conditions, the red outline becomes green after the delay, cuing production. **B**: Stimuli consisted of high lexical frequency (Snodgrass^36^) pictures, low lexical frequency pictures, words, and pseudowords. **C:** A cohort of 20 healthy subjects completed the delayed naming paradigm. The linear mixed-effects model of response time as the dependent variable with subject as a random effect (using Type III ANOVA using Satterthwaite’s method) with demonstrated significant main effects of delay, F_2,3390.6_ = 921.06, p < .001, and stimulus type, F_3,3388.7_ = 36.19, p < .001. There was also a significant interaction between delay and stimulus type, F_6,3387.2_ = 6.34, p < .001. 15 subjects who underwent intracranial electrophysiology with stereo EEG for drug-resistant epilepsy completed the paradigm. Similarly, there were significant main effects of delay, F_2,3176.6_ = 1087.78, p < .001, and stimulus type, F_3,3176.1_ = 79.64, p < .001 on response time. The interaction between delay and stimulus type was also significant, F_6,3176.0_ = 2.49, p = .021. **D-F:** Additional effects in the intracranial electrophysiology group. **D:** There were main effects of delay (F_2,3182.6_ = 1022.55, p < .001) and phonological density (F_1,3182.2_ = 79.86, p < 0.001) on response time. However, the interaction was not significant (F_2,3182.6_ = 0.92, p = 0.40), indicating that phonological encoding was not modulated by the delay period. **E:** Similarly, the main effect of single vs. multiple syllables was significant (F_1,3182.1_ = 38.62, p < 0.001), but the interaction between delay and number of syllables was not (F_2,3182.3_ = 1.82, p = 0.16). **F:** Finally, the main effect of repetition was significant (F_1,3190.0_ = 32.81, p < 0.001), but again the interaction between delay and repetition was not (F_2,3182.2_ = 0.31, p = 0.73).

### Psycholinguistic analyses

The primary psycholinguistic manipulation was stimulus type (high frequency picture, low frequency picture, word, pseudoword), which varied primarily in lexical frequency and visual recognition. Thus, we hypothesized that the delay would decrease response time after the go cue and that the effect of stimulus type would interact and diminish with delay. This primary analysis was confirmed using linear mixed effects modeling of response time (RT) with subject as a random effect in both healthy controls (delay: F_2,90_._6_ = 921.06, p < 0.001; stimulus type: F, _88_._7_ = 36.19, p < 0.001; delay x stimulus type, F_6,87_._2_ = 6.34, p < 0.001) and intracranial electrophysiology subjects (delay: *F*_2,176_._6_ = 1087.78, *p* < 0.001, stimulus type: *F*, _176_._1_ = 79.64, *p* < .001; delay x stimulus type: *F*_6,176_._0_ = 2.49, *p* = 0.021; **Figure 1C**). This behavioral finding indicates that processes sensitive to stimulus type (e.g. visual recognition and lexical selection) occur before the delay, permitting them to proceed or reach completion prior to the go cue in delayed conditions.

In contrast, other manipulations did not show interaction with delay. Phonological density of the stimuli (e.g the number of phonological neighbors a given word has) is a manipulation of phonological encoding^33^, with higher density words expected to produced faster than low. Should phonological encoding occur on demand when speech is cued, the effect of phonological density should not interact with delay. We observed a main effect of phonological density (*F*_1,182_._2_ = 79.86, *p* < 0.001), but the interaction of density with delay was not significant (*F*_2,182_._6_ = 0.92, *p* = 0.40; **Figure 1D**). Similar main effects but lack of interaction with delay were seen for number of syllables (main: *F*_1,182_._1_ = 38.62, p < 0.001; interaction: *F*_2,182_. = 1.82, p = 0.16; **Figure 1E**) and repetition (main: *F*_1,190_._0_ = 32.81, p < 0.001; interaction: *F*_2,182_._2_ = 0.31, p = 0.73; **Figure 1F**). Thus, we observe a functional dissociation between the effects of delay on visual recognition and lexical selection vs its lack of effect on syllable planning and phonological encoding.

### Electrode coverage

The twenty intracranial electrophysiology patients underwent stereo EEG implantation for workup of their drug-resistant epilepsy (**Figure 2A, Supplementary Figure 1**). While the primary purpose of the implantation was to interrogate one or more hypotheses about their epileptogenic zone, these implantations provided broad coverage of the lateral and medial surfaces of the cerebrum and deep structures including sulci, the insula, and subcortical structures, with densest coverage over the left perisylvian and basal temporal regions (**Figure 2B**).

**Figure 2:**
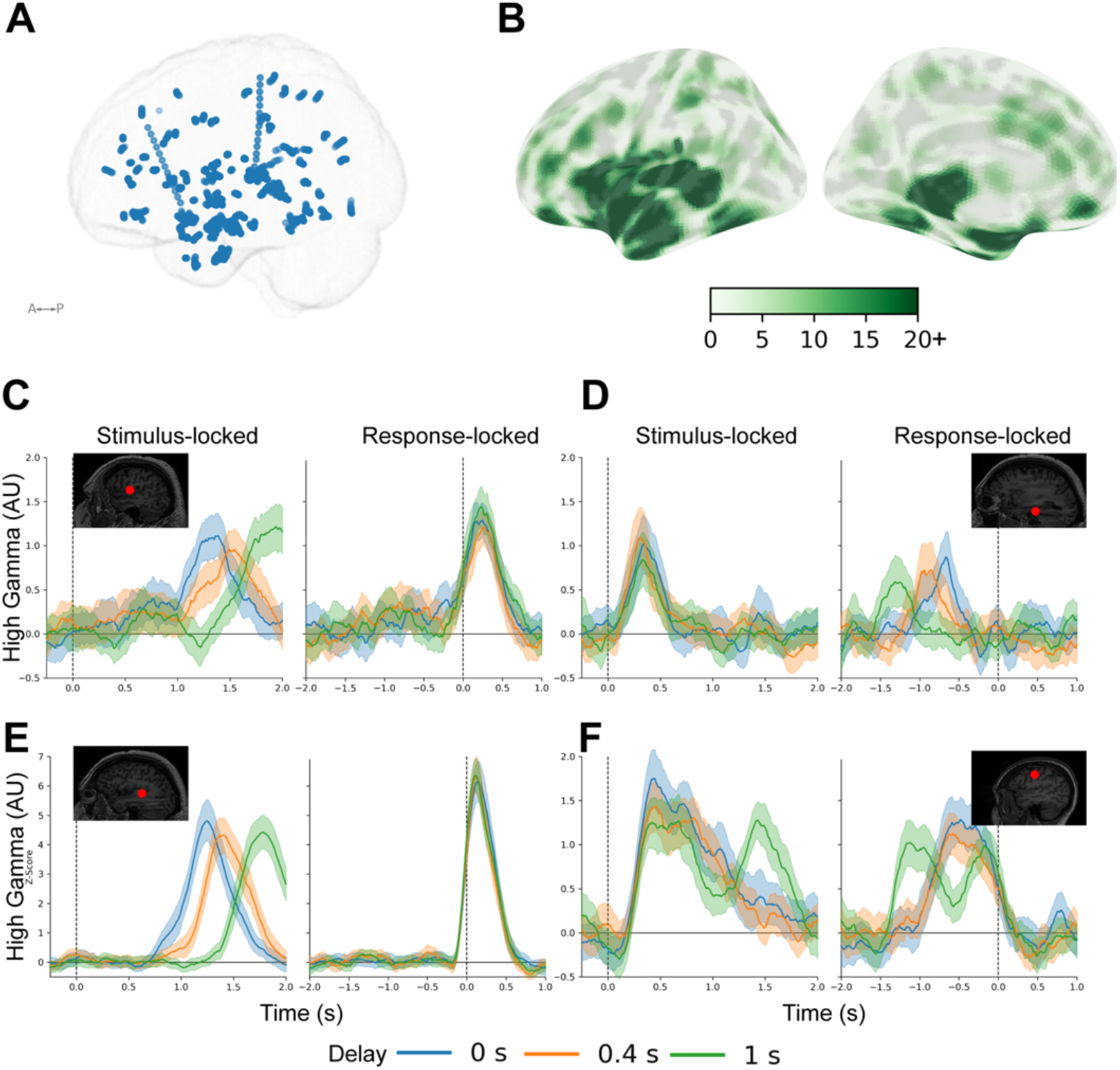
Electrode distribution and activity. **A:** Electrode contact distribution across 15 subjects. Electrodes were placed primarily to interrogate specific hypotheses about the subject’s epileptogenic zone but secondarily permitted broad exploration of both medial and lateral cortical surfaces and subcortical structures. Electrode shafts were primarily placed orthogonal to the lateral surface and consist of 8 - 18 electrode contacts number medial to lateral. **B:** Electrode contact density map was determined by counting the number of contacts within a 1cm radius. There was dense coverage of the perisylvian and basal temporal regions as well as broadly across the frontal and parietal lobes including the medial surface. **C-F**: Broadband gamma power (70-200 Hz, Z-scored to baseline) across multiple contacts showing typical patterns. Insets: electrode contact location on subject MRI in native space. Trials were epoched in stimulus-locked (aligned to the stimulus onset at 0s) and response-locked (aligned to the onset of speech at 0s). Specific contacts demonstrated differential alignment in each type of locking. Blue: No (0s) delay, Orange: 0.4s delay, Green: 1s delay. The solid line is mean activation across all trials with that delay while shading represents standard error of the mean. **C:** Speech motor cortex demonstrates strong alignment to response across all delay conditions. **D:** Contact in basal temporo-occipital cortex demonstrates strong stimulus-locking. **E:** Contact in primary auditory cortex in Heschl’s gyrus demonstrates extremely strong locking to the response (auditory feedback to hearing one’s own voice). **F:** Some contacts demonstrate alignment to both the stimulus and response. Perhaps surprisingly this contact in the junction of inferior frontal sulcus and precentral sulcus demonstrates strong locking with stimulus and response.

### Electrode responses

The high gamma (70-200 Hz) amplitude was extracted from the local field potential (LFP) at each contact and Z-scored relative to a pre-stimulus baseline (500ms before stimulus). Electrophysiological data was divided into epochs aligned either to the stimulus at time 0 s (stimulus-locked) or response at time 0 s (response-locked). Averaging over trials after epoching identifies electrodes that are more aligned to the response (**Figure 2C, E**) or stimulus (**Figure 2D**). For instance, an electrode contact in speech motor cortex (**2C**) showed variability in the timing of activation by delay in stimulus-locked but showed high alignment to response onset in response-locked epoching. In contrast, an electrode in basal temporo-occipital cortex (**2D**) demonstrating strong stimulus-locking. Primary auditory cortex in Heschl’s gyrus showed strong response-locking (**2E**), consistent with its role in self-monitoring of speech. Interestingly, many electrodes demonstrated intermediate stimulus/response locking or locking to both. For instance, an electrode at the junction of inferior frontal sulcus (IFS) and precentral sulcus (PreCS) demonstrated strong locking to stimulus at the start of its peak and response at the end (**Figure 2F**).

### Stimulus-response correlation space

We sought to quantify the degree of stimulus/response-locking for each contact using a noise-invariant approach. For each electrode contact we computed pairwise trial to trial correlation (within subject) and averaged over all pairs. This stimulus/response correlation captures the degree to which a given contact’s response aligns to the stimulus or initiation of response. This approach demonstrated high stimulus correlation in regions sensitive to stimulus (e.g. basal temporal occipital and parietal), high response correlation in regions sensitive to response (speech motor cortex, supplementary motor area, anterior temporal lobe), and several regions with sensitivity to both (**Figure 3A**).

**Figure 3:**
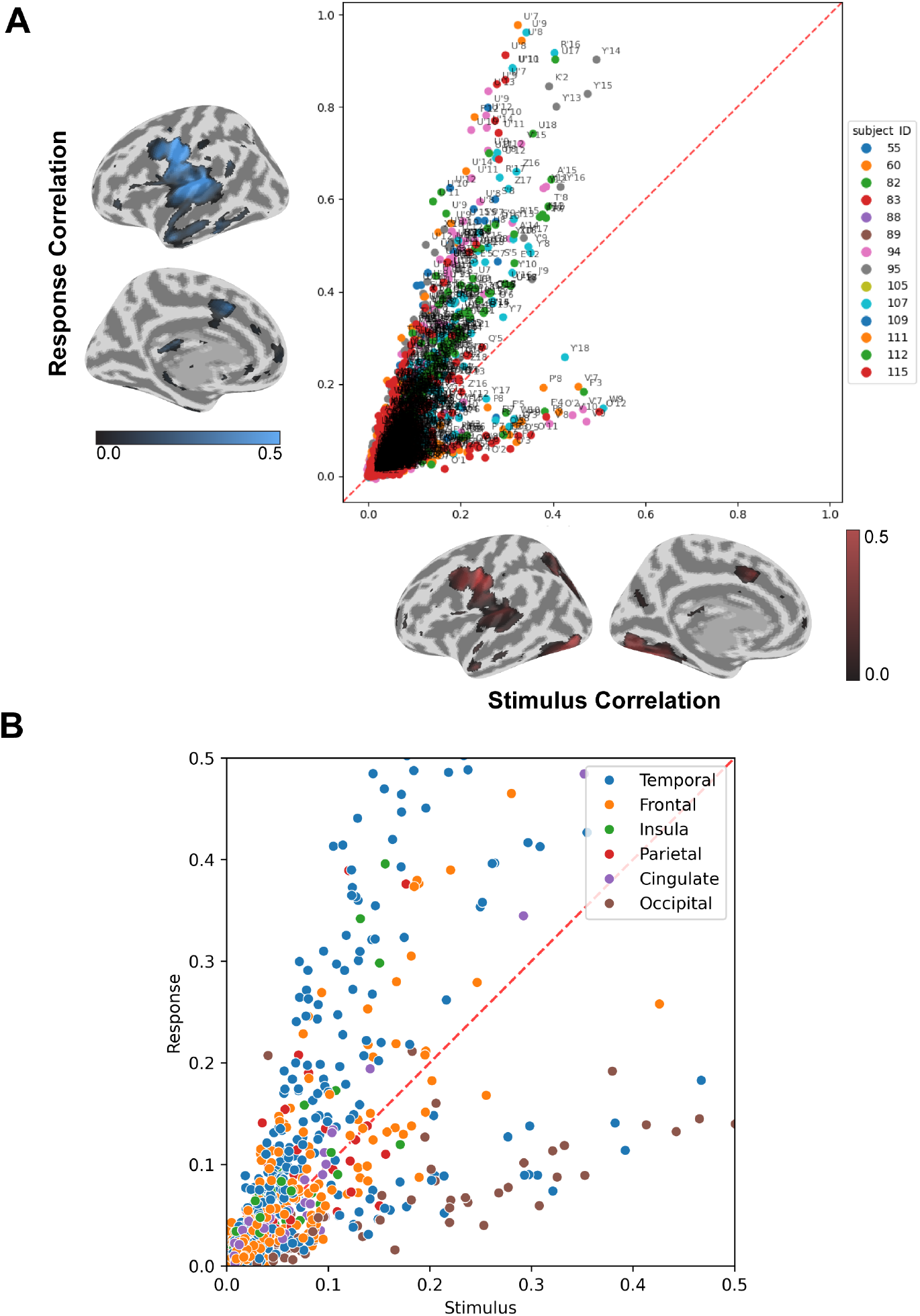
Stimulus-response correlation space. To quantify the degree of stimulus vs. response locking we calculated the trial-to-trial correlation across trials for each electrode in stimulus epoching and response epoching. If correlation in each epoching was high, this suggested that electrode was highly time-locked to stimulus or response. A: The scatterplot demonstrates the location of each contact in stimulus-response correlation space. Certain electrodes, such as those in speech motor cortex (R’16) or primary auditory cortex (U’7-10) demonstrated strong response correlation while others such as O’12 in the occipital lobe showed stimulus correlation. Note that each subject has contacts in various parts of correlation space. Cortical plots: Distribution of stimulus and response correlation along the cortex. B: Contact location by lobe demonstrates a broad distribution in correlation space. While the occipital lobe contacts are largely confined the stimulus-correlation half, most lobes (e.g. temporal and frontal) had contacts in both response and stimulus locked regions. Euclidean distance between contacts showed poor correlation to distance in correlation space (R^2^ = 0.013) suggesting that physical distance was irrelevant for position in correlation space.

Using this approach, we mapped all contacts over all subjects into a common stimulus-response correlation space (**Figure 3A**). In this space, contacts with response correlation greater than stimulus correlation are above the diagonal, while those with the opposite are below. At a global level we identified a bilobed structure, seemingly with limits at the extremes of response and stimulus correlation. Each lobe corresponds to a small subset of contacts with high response or stimulus correlation, but we note that the vast majority of electrodes show sensitivity to both (near the diagonal).

Other than contacts in the occipital lobe, location of contacts by lobe did not cluster in a particular region in correlation space (**Figure 3B**). While there is some common distribution of certain cortical regions (Harvard-Oxford Atlas, **Supplementary Figure 2**), there are multiple regions that span over several parts of correlation space. Euclidean distance in physical space between contacts showed poor correlation with distances in correlation space (R^2^ = 0.013). Collectively, this suggests that position in correlation space represents an independent dimension of cortical function, dependent entirely on the temporal properties of activation in each subregion.

### Psycholinguistic manipulations create shifts in stimulus-response correlation space that identify cortical regions sensitive to linguistic properties

In the previous analysis, trial-to-trial correlations were conducted at each electrode over *all* trials for a given subject. We subsequently sought to explore the effects of psycholinguistic manipulations (e.g. stimulus type, lexical frequency, phonological density) on correlation space. Thus, we calculated the trial-to-trial correlations at each contact for each type and level of manipulation (e.g high frequency vs. low frequency pictures).

At a global level, there seemed to few changes in the underlying bilobed structure of stimulus-response correlation space (**Supplementary Figure 3**). However, at the level of individual contacts, there were shifts in the stimulus and response correlations (**Figure 4, Supplementary Figure 4**). After thresholding for a minimum shift of 0.025, we classified contacts by the sign of change in stimulus correlation and response correlation, generating 4 clusters – 1: (-, -), 2: (-, +), 3: (+, -), 4: (+,+). Visualizing these contacts on the cortical surface demonstrated specific distributed networks with similar correlational shifts for each manipulation. For instance, for high vs. low frequency pictures, cluster 3 (+, -) consisted of a cluster in posterior middle temporal gyrus, planum temporale, and parietal lobe, while cluster 4 (+, +) included these regions as well as IFS/PreCS junction and insula (**Figure 4A**). Temporally, the mean high gamma activation within clusters was highly consistent and visually demonstrated sensitivity in amplitude and peak time to lexical frequency. We observed similar results for phonological density, with cluster 4 (+, +) identifying regions near pars triangularis, inferior frontal sulcus, and speech motor cortex and with cluster 1 (-, -) showing strong temporal variation between high and low phonological density and spatially spanning a large cortical distribution (**Figure 4B**). Similar results can be seen for pictures vs. text, words vs. pseudowords, and multi vs. single syllable words (**Supplementary Figure 4**).

**Figure 4:**
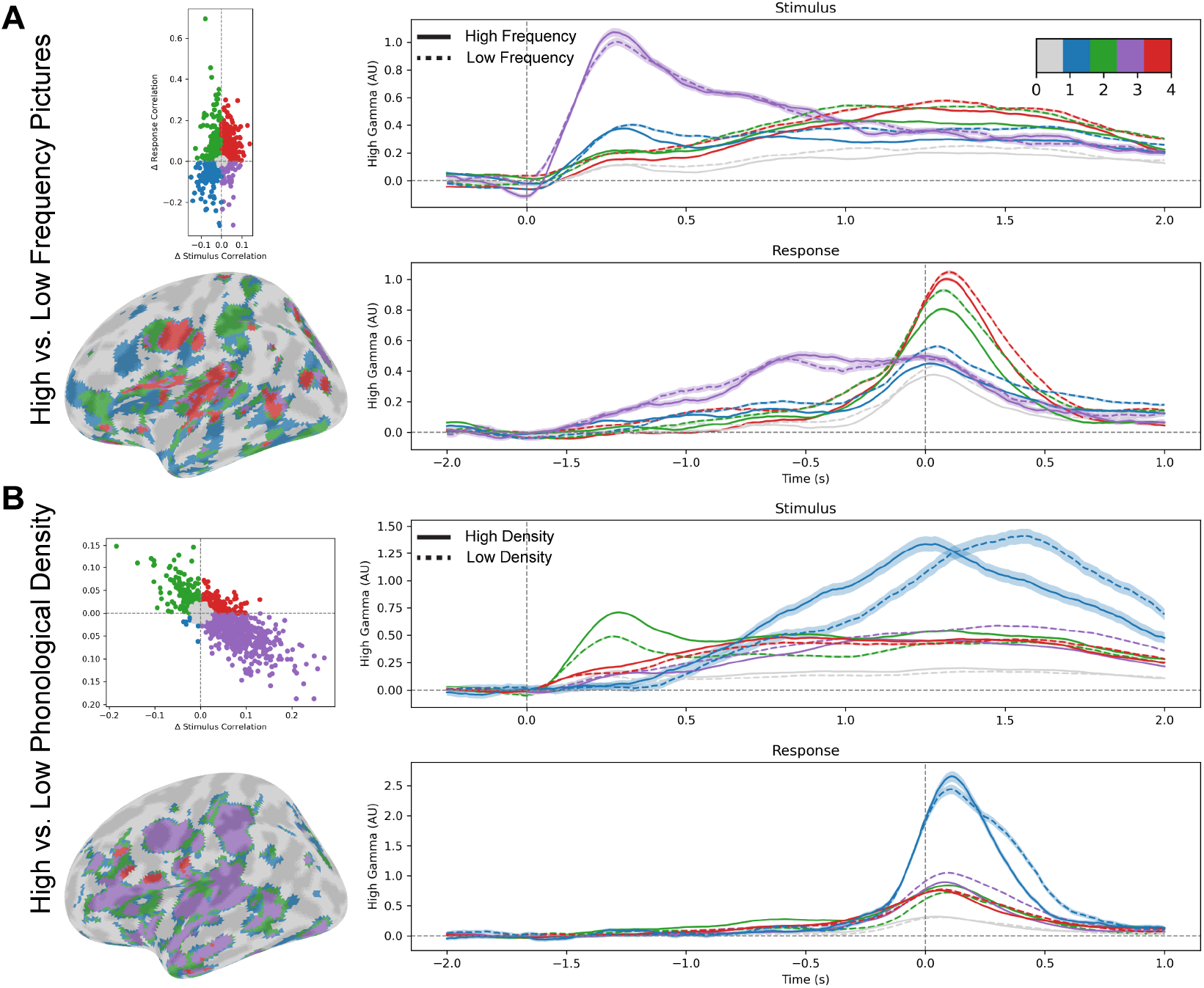
Psycholinguistic manipulations induce shifts in stimulus-response correlation space. Trial-to-trial correlation for each electrode conducted on specific subsets of all trials (e.g high frequency pictures vs. low frequency pictures) electrode contact location shifted in correlation space (top right). Electrodes were thresholded at a minimum shift of 0.025 then classified into the direction of the shift in stimulus-response space – e.g. Δ stimulus correlation, Δ response correlation (-,-), (-, +), (+, -), (+,+). This classification over change in stimulus-response correlation space corresponded with a cortical segmentation (bottom right) of regions showing differential responses to a given psycholinguistic manipulation (e.g lexical frequency, phonological density). Average high gamma power over contacts in a given class across all subjects showed specific and different patterns of activation to the psycholinguistic manipulation (right). Shifts in correlation space along each axis were correlated against changes in response time. A: Lexical frequency. Cortical regions including the posterior MTG and STS, lateral occipital lobe, IFS/PreCS, insula, and other regions demonstrated specific patterns of response. Δ Stimulus correlation in class 3 (purple) explained 34% of variance in response time between high and low frequency picture conditions by subject, while Δ response correlation in class 1 (blue) explained 31%. B: Phonological density. A broad swath of cortical regions demonstrates differential effects to phonological density. Shift in stimulus correlation in Class 4 (red) regions, consisting of inferior frontal sulcus, pars triangularis, the anterior ascending ramus of the Sylvian fissure and speech motor cortex explained 32% of variance in response time between high and low phonological density stimuli across all subjects, while shift in stimulus correlation in class 1 (blue) explained 49%. Additional psycholinguistic manipulations (pictures vs. text, pseudowords vs. words, multiple vs. single syllables) can be seen in Supplementary Figure 4.

### Variance in response time is explained by shifts in stimulus-response correlation space

We examined if shifts in correlation space induced by a given psycholinguistic manipulation corresponded to variance in RT from that manipulation at the subject level. For instance, we computed the mean change in stimulus correlation and response correlation for each cluster and regressed this with difference in RT between high and low lexical frequency pictures treating each subject as the unit of observation. For lexical frequency, variance in stimulus correlation shifts in cluster 3 (+, -) explained 33.9% of the variance in RT, response correlation shifts explained 27.3%, and together they explained 64.0% of variance in RT. For phonological density, linear regression of cluster 1 (-, -) stimulus and response correlation shift against change in RT explained 74.5% of variance. For words vs. pseudowords, cluster 2 (-, +) explained 23.9% of variance in RT. Finally, for multiple vs. single syllables, cluster 2 explained 20.3% of variance in RT. Thus, the clustering imposed by shifts in correlation space created psycholinguistically meaningful clusters for each tested manipulation.

### Stimulus-response correlation space analysis identifies key steps in language production across subjects

Finally, we tested if in aggregate correlation shifts in all relevant cortical clusters could explain overall variability in trial level RT across all subjects. We regressed correlation shifts of all variables (stimulus type, picture lexical frequency, phonological density, number of syllables, repetition, and delay) for each subject against trial level RT across all subjects. We used elastic net linear regression with L1 and L2 regularization (regularization parameter = 2, L1 ratio = 0.8) to account for multicollinearity, overfitting, and model selection with leave-one-subject-out cross-validation across all 15 subjects. This regression approach demonstrated that shifts in correlation space induced by these manipulations could explain 31.4% of variance in trial level RT.

Finally, we plotted activity in individual clusters for each psycholinguistic variable to understand the spatiotemporal distribution of specific psycholinguistic processes in the brain (**Figure 5**). We identified specific processes sensitive to each variable – e.g. lexical selection to picture lexical frequency or phonological encoding to phonological density. The resulting patterns of activity corroborated our behavioral observations. For picture naming in the immediate (0 ms) vs. delayed (1000 ms) conditions, lexical selection peaked at ∼250ms in both conditions, while syllable planning and phonological encoding shifted as a function of delay at the time of response (**Figure 5A**). For high vs. low frequency pictures, the amplitude of lexical selection was slightly smaller for low frequency pictures, and subsequent steps shifted later temporally, resulting in a frequency effect on RT (**Figure 5B**). For pictures vs. words, the amplitude of lexical selection was reduced for words and phonological encoding shifted earlier (**Figure 5C**). Finally for words vs. pseudowords, lexical selection was minimal for both conditions, but syllable planning and phonological encoding were later for pseudowords as expected. The cortical locations of these clusters can be seen in **Figure 5E-H** and **Supplementary Figure 4**. Thus, using a classification approach based on shifts in stimulus-response correlation space, we can identify the electrophysiological signatures and cortical localizations of each stage of linguistic processing, thereby providing a spatiotemporal account of language production.

**Figure 5:**
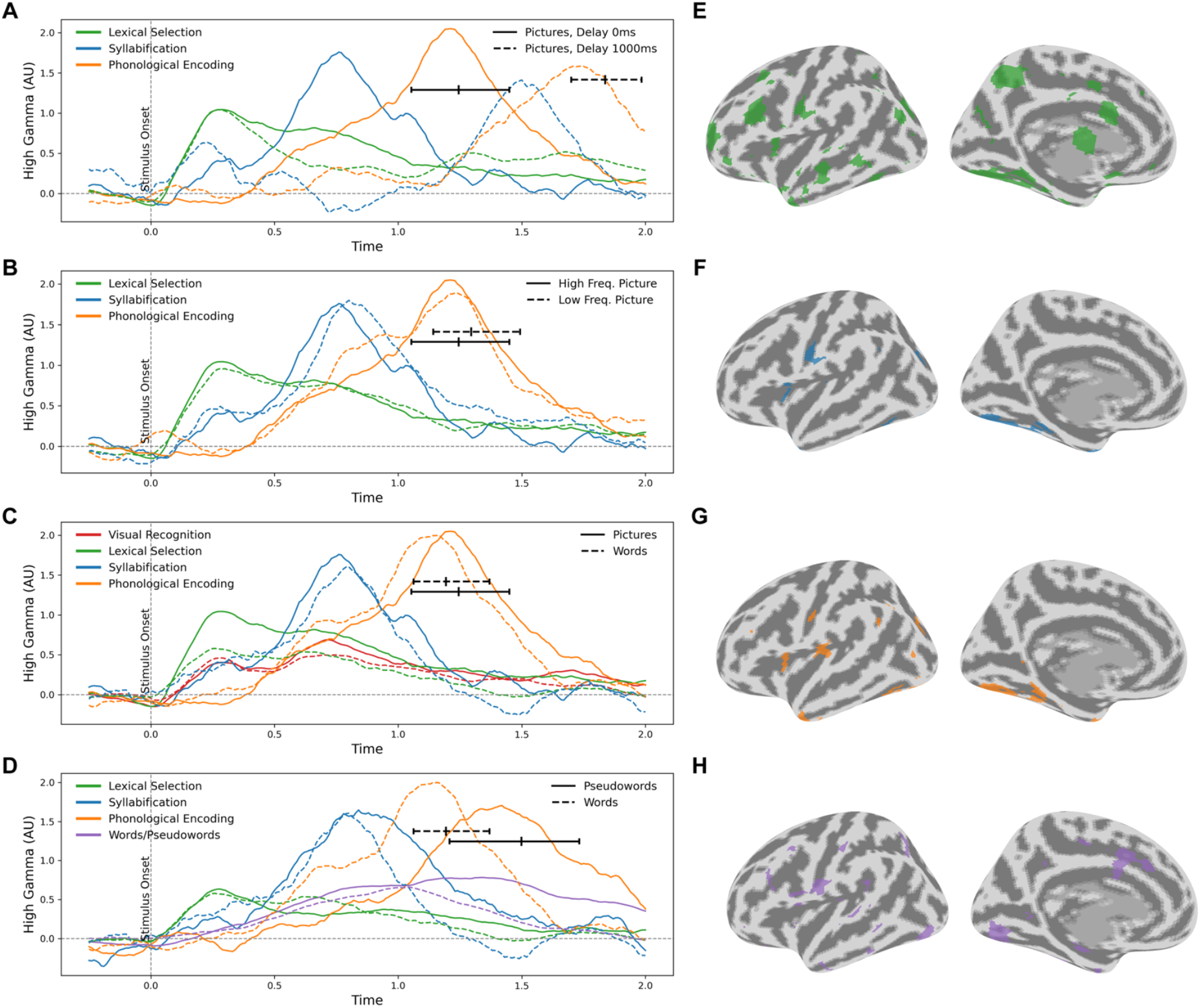
Change in correlation space identifies cortical networks responsible for specific aspects of language production. Using leave-one-subject-out cross-validation with linear regression with L1 and L2 regularization (e.g. elastic net regularization, regularization weight = 2, L1 ratio = 0.8) identified cortical networks (classes) that in aggregate best predicted overall variance in response time across subjects and collectively explained 31.4% of variance in response time. Networks demonstrating specific sensitivities to psycholinguistic manipulations are demonstrated. Black lines with ticks correspond to the 25^th^, 50^th^, and 75^th^ percentiles of mid-speech duration by condition. **A:** High frequency pictures between the 0 ms and 1000 ms delay conditions. While the network corresponding to lexical selection shows shifts in amplitude, the overall temporal distribution is unaffected by delay. In contrast, syllable planning and phonological encoding shift with the delay, corresponding to shifts in response time. **B:** High vs. low frequency pictures (0s delay) demonstrate similar patterns of activation in lexical selection, but syllable planning and phonological encoding demonstrates a shift to later timepoints corresponding with the longer response time. **C:** Pictures vs. words (0s delay). Cortical regions showing sensitivity to visual recognition demonstrate similar activation pattern between pictures and words, but those corresponding to lexical selection show less activation for words. Phonological encoding also occurs earlier for words than pictures. **D:** Pseudowords vs. words (0s delay). Overall activation in lexical selection regions is low. Syllabification and phonological encoding are delayed in pseudowords. A broad network demonstrating shifts in correlation space with words vs. pseudowords also demonstrates broadly greater activation for pseudowords over words. **E-H: Spatial maps of cortical networks underlying specific psycholinguistic stages. E:** Lexical selection. **F:** Syllable planning. **G:** Phonological encoding. **H:** Words vs. pseudowords.

## Discussion

Using a delayed naming paradigm with specific psycholinguistic manipulations in conjunction with stimulus-response correlation space analysis of intracranial electrophysiology, we identify specific cortical networks underlying specific stages of language production. With the response time behavioral analysis in healthy and intracranial electrophysiology subjects we confirm our hypothesis that some effects precede the delay period and are correlated to stimulus onset, e.g. lexical access. Given that the delay permits these functions to complete, their effect diminishes (interacts with) the delay period with respect to RT. In contrast, other manipulations, such as phonological density, are not modulated by delay and occur on demand once speech production is cued. Thus, the behavioral results support a model with discrete stages of speech processing.

Stimulus-response correlation space analysis exploits temporal variance in response time with the delay and variation in stimulus type, lexical frequency, phonological density, etc. and determines the degree to which electrophysiological data is aligned to stimulus or response (or intermediate stages). This provides a novel methodology for analyzing data across different implantations and subjects without restricting analysis to specific cortical regions, a common issue in intracranial electrophysiology studies^46^. Stimulus-response correlation space has a seemingly invariant structure at the global level, with one population of contacts showing strong response correlation, another showing strong stimulus correlation, and the vast majority of contacts showing an intermediate degree. This is concordant with views of cortical organization that identify specific sensory and effector regions bridged by broad regions of association cortex^47–50^.

Despite a lack of change in the macroscopic structure of correlation space, specific psycholinguistic manipulations induce shifts of electrode contacts and cortical regions in correlation space to higher/lower stimulus/response correlation. The conceptual interpretation of the direction of shift is challenging but likely represents a temporal shift in processing in a given cortical region with some contribution from amplitude shifts. By classifying electrodes and cortical regions by the direction of change with a given manipulation, we develop a parcellation of the cortex sensitive to this manipulation and thus involved in the underlying processing^1,2^. Correlation shifts in response to a psycholinguistic manipulation are highly predictive of variance in response time due to that manipulation – up to 64% of the variance due to phonological density and 74.5% of variance due to lexical access. We then validated this approach at predicting trial-level variance of response time across subjects, using leave-one-out cross validation approach, explaining 31.4% of variance. The cross-validation suggests that stimulus-response correlation space shifts have an invariant relationship to the temporal processing of speech. We acknowledge that this reduction in correlation could be due to incorrect cortical parcellation. However, this could also be due to unaccounted processes in language production in our study design. Collectively these results identify spatiotemporally separated cortical networks that underlie speech production that seemingly proceed in a temporally separated but overlapping sequence.

This approach builds on and validates prior investigations of specific stages of speech processing. We find regions in inferior frontal gyrus/frontal operculum and speech motor cortex sensitive to phonological planning and syllable production, similar to prior accounts^51,52^. Regions sensitive to lexical access are found broadly across the cortex, in contrast to prior reports identifying the posterior MTG primarily^53,54^. In particular, regions not typically explored with EEG or subdural grid electrocorticography, including the inferior frontal sulcus and supplementary motor area, were implicated. Indeed, the role of the depths of sulci and medial regions may be underappreciated when comparing our results to other noninvasive electrophysiological approaches.

Beyond great relevance for cognitive neuroscience, these results also have direct implications for clinical neurosurgery and neurosciences. An expanded neurocognitive model has direct implications for what can and should be considered “eloquent” in resective brain surgery and how this should be mapped^17,56^. The eloquence of each of these regions with respect to inducing a transient or permanent deficit will need to be validated. Furthermore, the existence of seemingly poly-functional regions – e.g. inferior frontal sulcus – may suggest that our parcellation of speech function may not map directly onto cortical processing^57^. The white matter pathways underlying these cortical networks is also of crucial importance and will be a focus of future investigation. Finally, other “network” pathologies of the brain such as epilepsy may exploit the same cortical and cortical-subcortical networks that underlie speech processing^13,58^ resulting in specific semiological features.

Finally, we acknowledge that high gamma activity represents only a subset of cortical and cortical-subcortical activity. Although high gamma is most closely tied to local neuronal firing and BOLD activation^59–61^, lower LFP frequencies in the alpha and beta range may represent cortico-cortical or cortico-subcortical top-down modulation^62^. This modulation in lower frequency ranges may better capture cascading aspects in speech production, highlighting the need for detailed neurophysiological analysis of brain activity. Furthermore, connectivity between regions may occur in the high gamma range or may be better represented with other frequency ranges and/or connectivity measures. Interactions between frequency bands and top-down modulation are of key interest in future work.

Stimulus-response correlation space analysis can examine the temporal shifts induced by specific psycholinguistic manipulations and identify the spatiotemporal signatures of language processing. This flexible tool may be applicable to other studies of language and cognition with specific temporal manipulations in the behavior study design. Using this approach, we identify a sequential but overlapping process of speech production that is distributed in broad networks across the cortex.

