## Supplemental Figures 1-4 for "Stimulus-response correlation analysis dissociates spatiotemporal cortical networks supporting speech production"

**Supplementary Figures:**

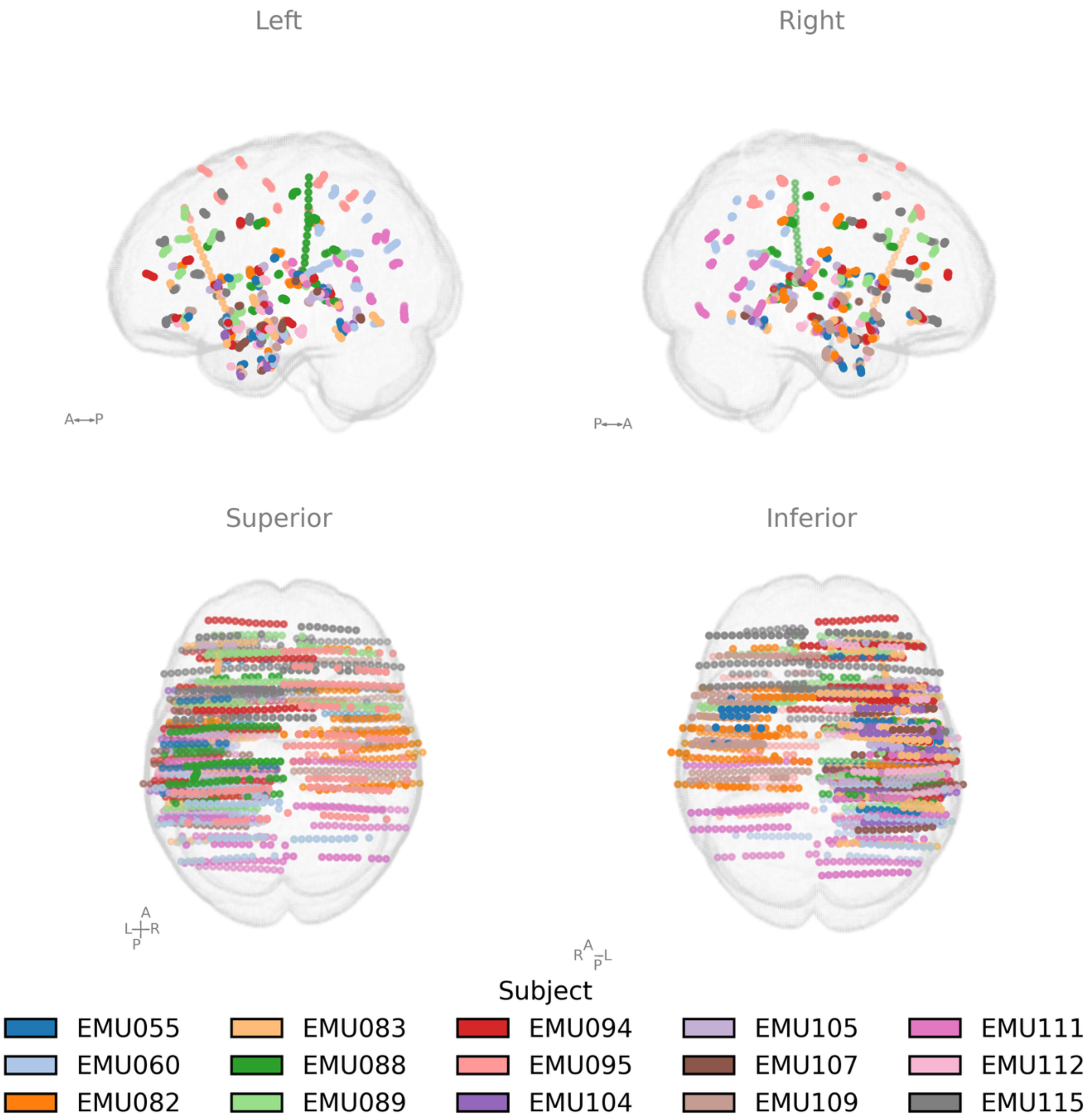


**Supplementary Figure 1:** Electrode coverage by subject demonstrates broad coverage of both hemispheres of both dorsolateral and medial cortical surfaces as well as subcortical structures.

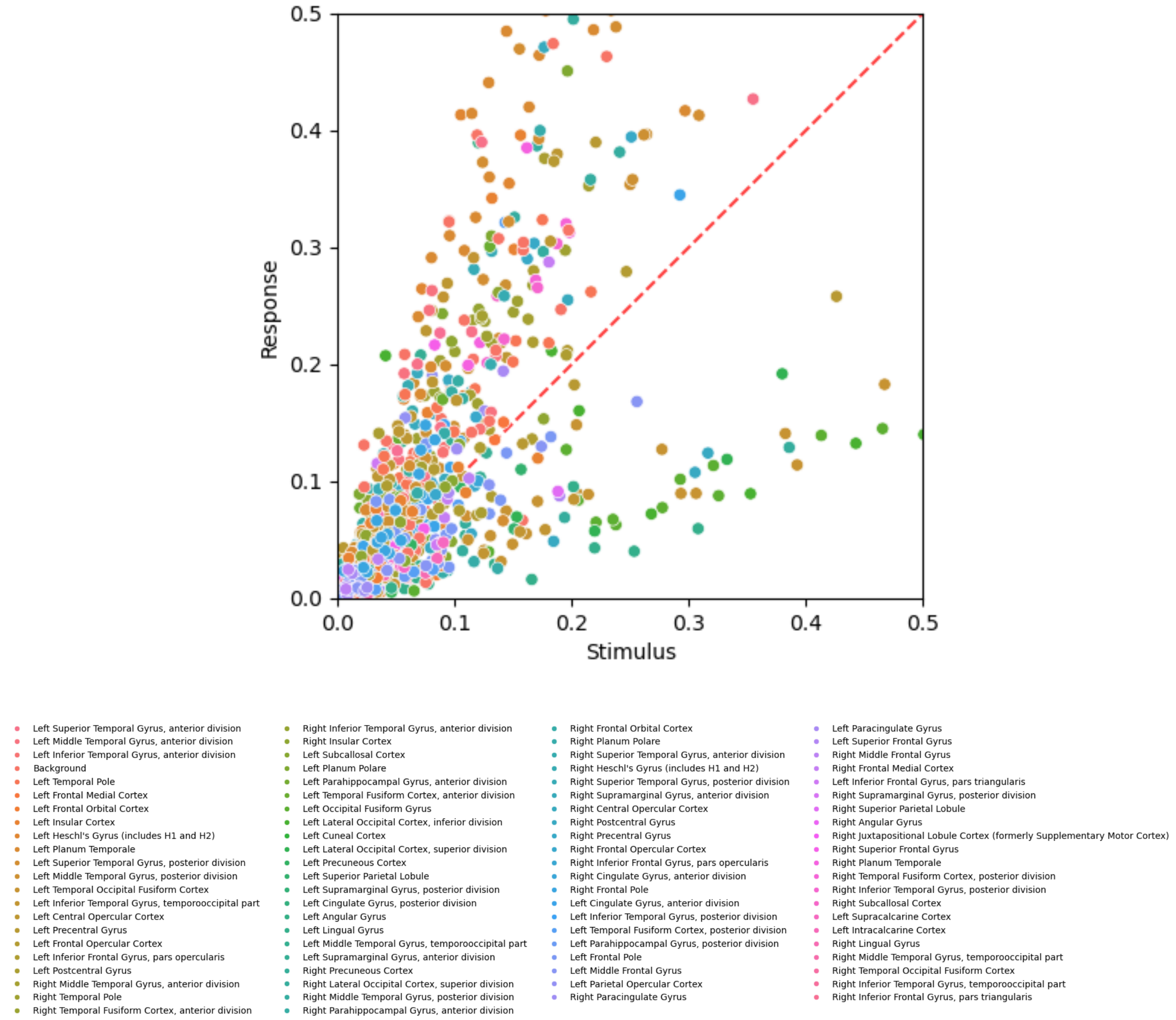


**Supplementary Figure 2: Stimulus-response correlation space vs. cortical subregions** (Harvard-Oxford Atlas). While specific cortical regions (e.g. occipital and basal temporal cortex) do demonstrate specific clustering in correlation space, others show broad distribution.

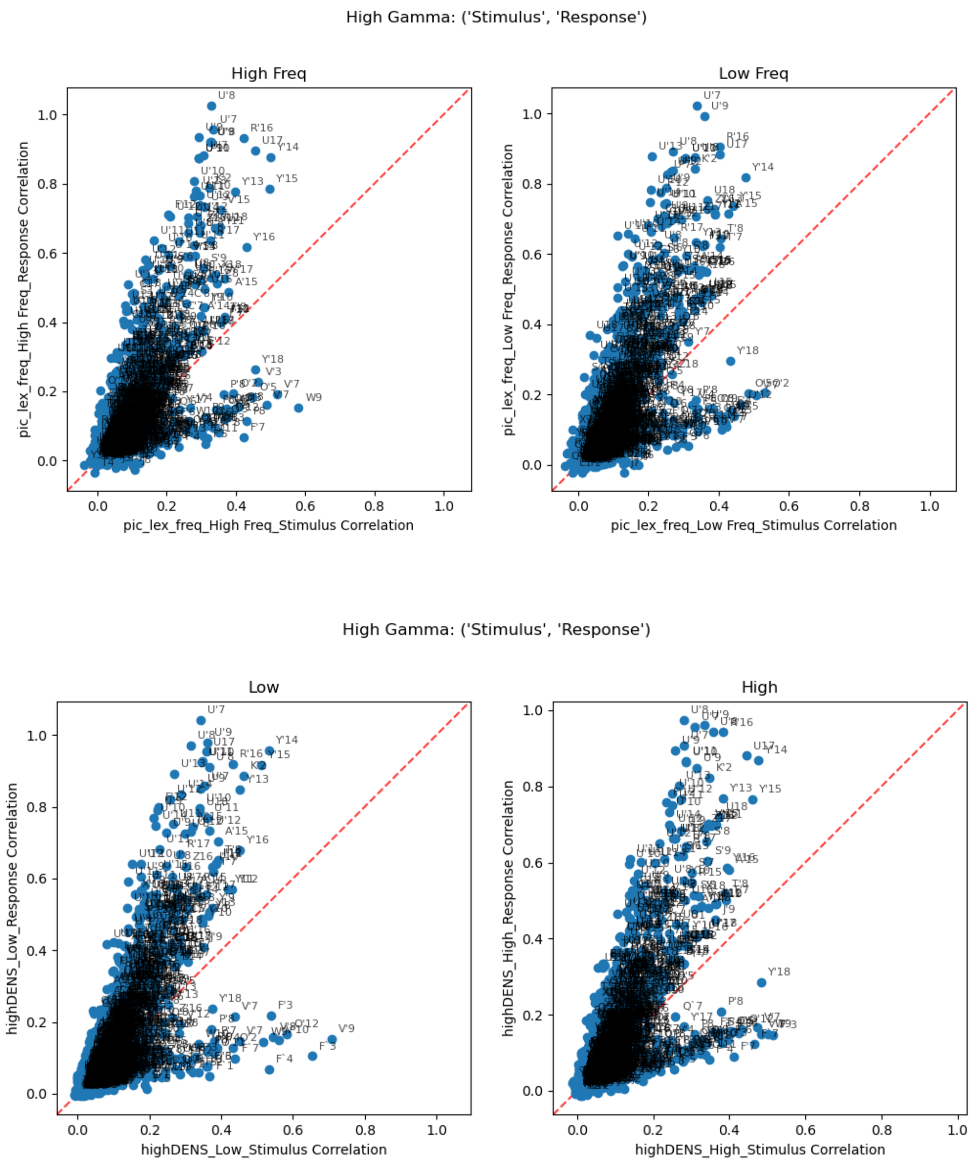


**Supplementary Figure 3:** Trial-to-trial correlation for individual electrode contacts across subjects across subsets of trials corresponding to picture lexical frequency (*top*), and phonological density *(bottom*. While there are shifts in correlation space of individual contacts, the global structure of stimulus-response correlation space seems invariant to specific psycholinguistic manipulations.

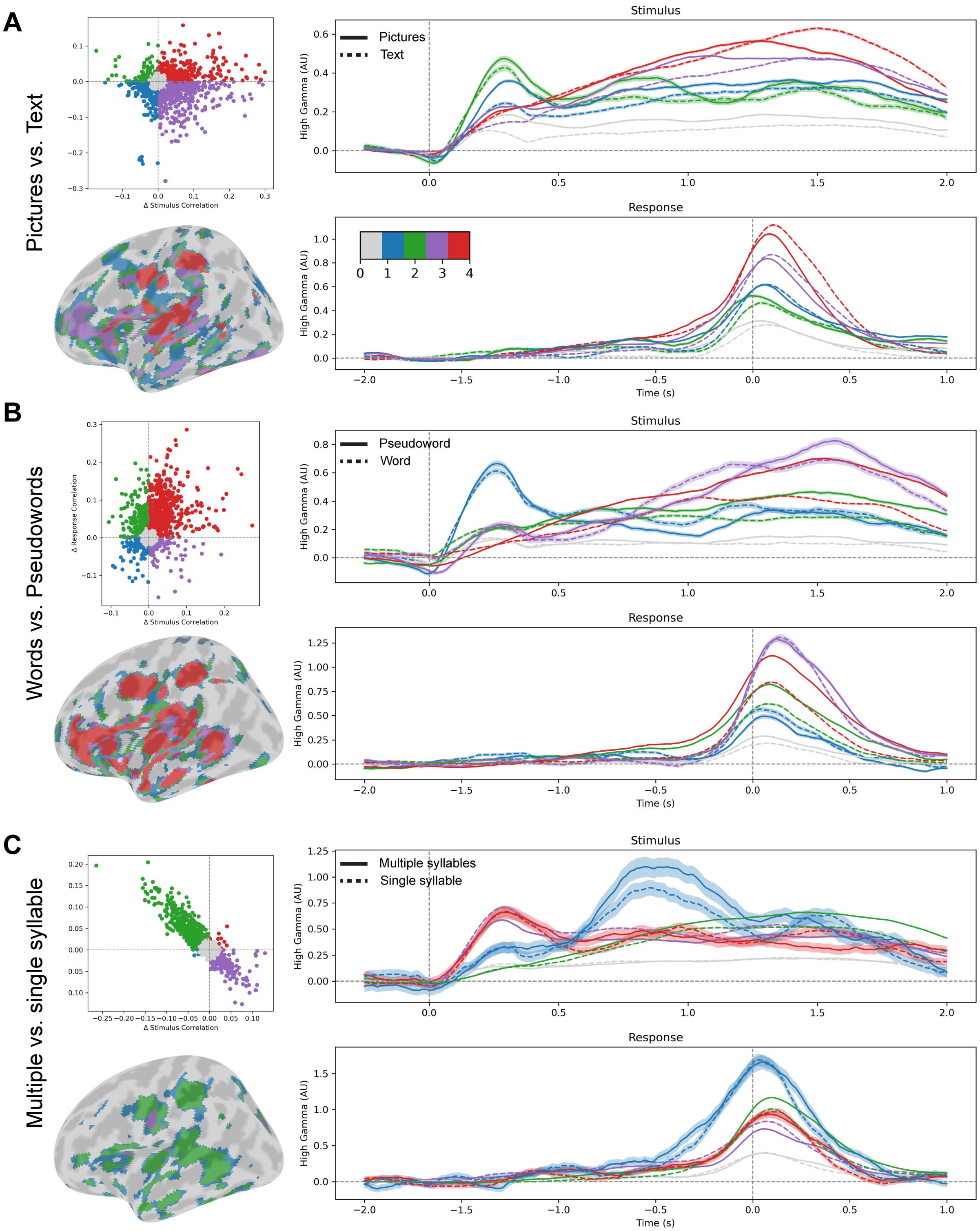


**Supplementary Figure 4:** Classification of electrode contacts and cortical regions by shifts in stimulus-response correlation space with psycholinguistic manipulations. **A:** pictures vs. text. **B:** words vs. pseudowords. **C:** Multiple vs. single syllable words.
